# Accuracy and Reliability of a Suite of Digital Measures of Walking Generated Using a Wrist-worn Sensor: Performance Characterization Study

**DOI:** 10.1101/2023.04.21.537226

**Authors:** Nathan Kowahl, Sooyoon Shin, Poulami Barman, Erin Rainaldi, Sara Popham, Ritu Kapur

## Abstract

**Background/Objectives:** Mobility is a meaningful aspect of an individual’s health whose quantification can provide clinical insights. Wearable sensor technology can quantify walking behaviors (a key aspect of mobility) through continuous passive monitoring. Our objective was to characterize the accuracy and reliability performance of a suite of digital measures of walking behaviors, as critical aspects in the practical implementation of digital measures into clinical studies.

**Methods:** We collected data from a wrist-worn device (the Verily Study Watch) worn for multiple days by a cohort of volunteer participants without history of gait/walking impairment in a real world setting. Based on step measurements computed in 10-second epochs from sensor data, we generated individual daily aggregates (participant-days) to derive a suite of measures of walking: step count, walking bout duration, number of total walking bouts, number of long walking bouts, number of short walking bouts, peak 30-minute walking cadence, peak 30-minute walking pace. To characterize accuracy of the measures, we examined agreement with truth labels generated by a concurrent, ankle-worn, reference device (Modus StepWatch 4™) with known low error, calculating the following metrics: Intraclass Correlation Coefficient (ICC), Pearson R, Mean Error (ME), Mean Absolute Error (MAE). To characterize the reliability, we developed a novel approach to identify the time to reach a reliable readout (time-to-reliability) for each measure. This was accomplished by computing mean values over aggregation scopes ranging from 1-30 days, and analyzing test-retest reliability based on ICCs between adjacent (non-overlapping) time windows for each measure.

**Results:** In the accuracy characterization, we collected data for a total of 162 participant-days from a testing cohort (N=35 participants; median observation time, 5 days). Agreement with the reference device-based readouts in the testing subcohort (n=35) for the eight measurements under evaluation, as reflected by ICCs, ranged between 0.7-0.9; Pearson R values were all greater than 0.75. For the time-to-reliability characterization, we collected data for a total of 15,120 participant days (overall cohort N=234; median observation time, 119 days). Here, all digital measures achieved an ICC between adjacent readouts > 0.75 by 16 days of wear time.

**Conclusions:** We characterized accuracy and reliability of a suite of digital measures that provides comprehensive information about walking behaviors in real-world settings. These results, which report the level of agreement with high-accuracy reference labels and the time duration required to establish reliable measure readouts, can guide practical implementation of these measures into clinical studies. Well-characterized tools to quantify walking behaviors in research contexts can provide valuable clinical information about general population cohorts and patients with specific conditions.

## Introduction

Assessing an individual’s mobility can provide meaningful insights into their general health status. In clinical settings, mobility is a fundamental factor to define prognosis and care as it is closely associated with a wide array of health outcomes [1–3]. However, accurate and reliable quantification of mobility in real-world settings remains challenging, since self-reported data from instruments such as the International Physical Activity Questionnaire can be biased by limited recall and social desirability [4,5].

The interest in quantifying physical activity using wearable devices has recently increased, as these technologies can collect objective, individualized data [6]. Wearable sensors have been incorporated into clinical studies across different disease states to enable movement analyses and the quantification of discrete physical activities, in order to develop clinically meaningful endpoints [7,8].

Yet, to cement their research utility, two aspects of these digital measurements need to be properly characterized: 1) the accuracy with which a digital measurement reads the parameters of interest [9], and 2) the amount of aggregated data needed to reliably capture an individual’s underlying behavioral state, minimizing noise related to natural variability, which usually translates into an aggregation time period for data collection (time-to-reliability). While accuracy is always a critical aspect in the characterization of a measure’s performance, time-to-reliability tends to be ignored, even though it is key for establishing fundamental study design specifications (e.g., collection time periods, length of wear time per day), to define baselines, or to compute power calculations for the detection of intervention effects or other changes.

Studies characterizing the performance of digital walking measures often focus on step count. However, the literature around these studies shows considerable heterogeneity across designs and some notable limitations. First, analyses tend to rely on truth labels originated by participants’ self reports, short-term close monitoring [10–12], or from reference devices with suboptimal accuracy (mean absolute percentage error >20%) and/or with the same body placement as the investigational devices, which would bias agreement results [13]. Second, these studies are often conducted in artificial laboratory environments, which inherently limit behavior range and are susceptible to subjectivity, assessment bias and/or unreliability [14–16]. Third, reliability characterization for investigational digital measurements is often absent from studies, despite having been acknowledged as an important element for the validation of clinically important research metrics, such as patient reported outcomes [17]. Beyond step counts, there have been studies that have used other digital measures to generate clinical insights, but without full characterization of their performance [18–29].

In previous work, we developed an algorithm that accurately classifies ambulatory status from data collected from a wrist-worn device, characterizing its performance across diverse demographic groups in a real-world setting [30]. Further, results from a substudy of an interventional randomized phase-2 trial demonstrated that digital measures of physical activity (step count and ambulatory time) could be sensitive to treatment effect in patients with Lewy Body dementia [31].

Herein, we report on the development of a series of measures that can capture walking behavior comprehensively, characterizing their accuracy and reliability. These measures included: (1) step count, (2) walking bout duration, (3) number of total walking bouts, (4) number of long walking bouts, (5) number of short walking bouts, (6) peak 30-minute walking cadence, and (7) peak 30-minute walking pace. To characterize their accuracy, we compared the measure readouts generated from a study device to highly accurate truth labels from an ankle-worn reference device, in healthy volunteers. To characterize their reliability, we developed a novel approach to calculate the aggregated time required to reach a reliable readout (time-to-reliability) for each measure.

## Methods

### Study Participants

The study cohort (Pilot program study) included adult volunteer participants, recruited amongst Verily Life Sciences employees in two locations (South San Francisco, CA and Cambridge, MA), without specific selection criteria. Gender and age information was collected for the accuracy characterization (not for the reliability characterization). This study was determined to be exempt research that did not require institutional review board review.

### Devices

The Verily Study Watch was the study device. This is a wrist-worn smartwatch that records acceleration data via an onboard inertial measurement unit (IMU) with 30Hz 3-axis accelerometer.

For the accuracy characterization, the ankle-worn Modus StepWatch 4™, an FDA-listed device, was used as reference from which to obtain ground truth labels for step counts and other derived walking measures. This is a single-axis accelerometer device with the greatest accuracy relative to other wearable devices, compared to human counting in real-world and in-lab settings [24,32].

### Generation of Digital Measurements

We collected continuous, raw accelerometer sensor data from the study smartwatch, computing step counts for every 10-second, non-overlapping epoch (for additional information about the algorithm to determine step counts, see Supplement), and collected step counts from the reference device also in 10-second epochs. From the 10-second epoch-based step counts, other measures of walking were derived, applying the same computations to the step counts from both devices (summarized in Table 1). We report the measure readouts as daily aggregates for individual participants (ie, participant-days).

**Table 1.** Summary of walking measure definitions

| Daily Walking Measure | Definition |
| --- | --- |
| Step count | Summed number of steps per day |
| A walking bout was defined as a series of contiguous 10-s epochs containing $\geq 6$ steps each and lasting for $\geq 30$ s (ie, at least three epochs). Epochs were considered contiguous if they were not interrupted by $>20$ s (ie, by no more than two 10-s epochs). | |
| Number of walking bouts | Summed number of walking bouts per participant-day |
| Number of short walking bouts | Summed number of walking bouts lasting between $\geq 30$ s and $< 1$ minute, per participant-day |
| Number of long walking bouts | Summed number of walking bouts lasting $\geq 2$ minutes per participant-day |
| Walking bout duration, mean | Mean duration of daily walking bouts |
| Walking bout duration, SD | Standard deviation of the duration of daily walking bouts |
| Walking bout duration, 95th percentile | Highest bout duration below the top 5% longest bouts |
| Walking cadence was defined as the number of steps per unit of time (in this study, per second [steps/s]) |  |
| Peak 30-minute walking cadence | For each participant-day, average cadence for the 180 10-s epochs (ie, 30 minutes, not necessarily contiguous) with the highest cadence. |
| Peak 30 minute walking pace | For each participant-day, average pace (calculated from the cadence and estimated stride length based on gender and height) for the 180 10-sec epochs (ie, 30 minutes) with the highest cadence (namely, the daily peak 30-minute cadence); measured as meter/s. |

s=second; SD=standard deviation

## Analyses

### Accuracy characterization

For the characterization of accuracy, the observation period ran from June-December 2019. The overall analysis cohort (N=70) was split into two equal (n=35), training and testing groups (Appendix Fig. 1). Participants were required to wear the two devices, the smartwatch and the reference device, throughout waking hours for up to 10 days. Step counts were obtained for both the study and the reference devices, for as long as both devices had been worn simultaneously by each participant, and filtered for days with ≥8 hours of weartime and >100 steps. Each subsequent measure was derived based on step counts from each device (Table 1) and compared for agreement. Agreement was examined using the following metrics: Fisher’s Intraclass Correlation Coefficient (ICC) as the main metric; Pearson R coefficient; mean error (ME); mean absolute error (MAE). For each metric, we calculated 95% CIs by bootstrapping with 1000 resampling iterations to account for multiple days (generally 5) from a given participant. Additionally, to further characterize the degree of agreement and bias of each measure, we examined measurements and distributions between devices and Bland-Altman Plots with 95% Limits of Agreement (LoA).

**Figure 1.**
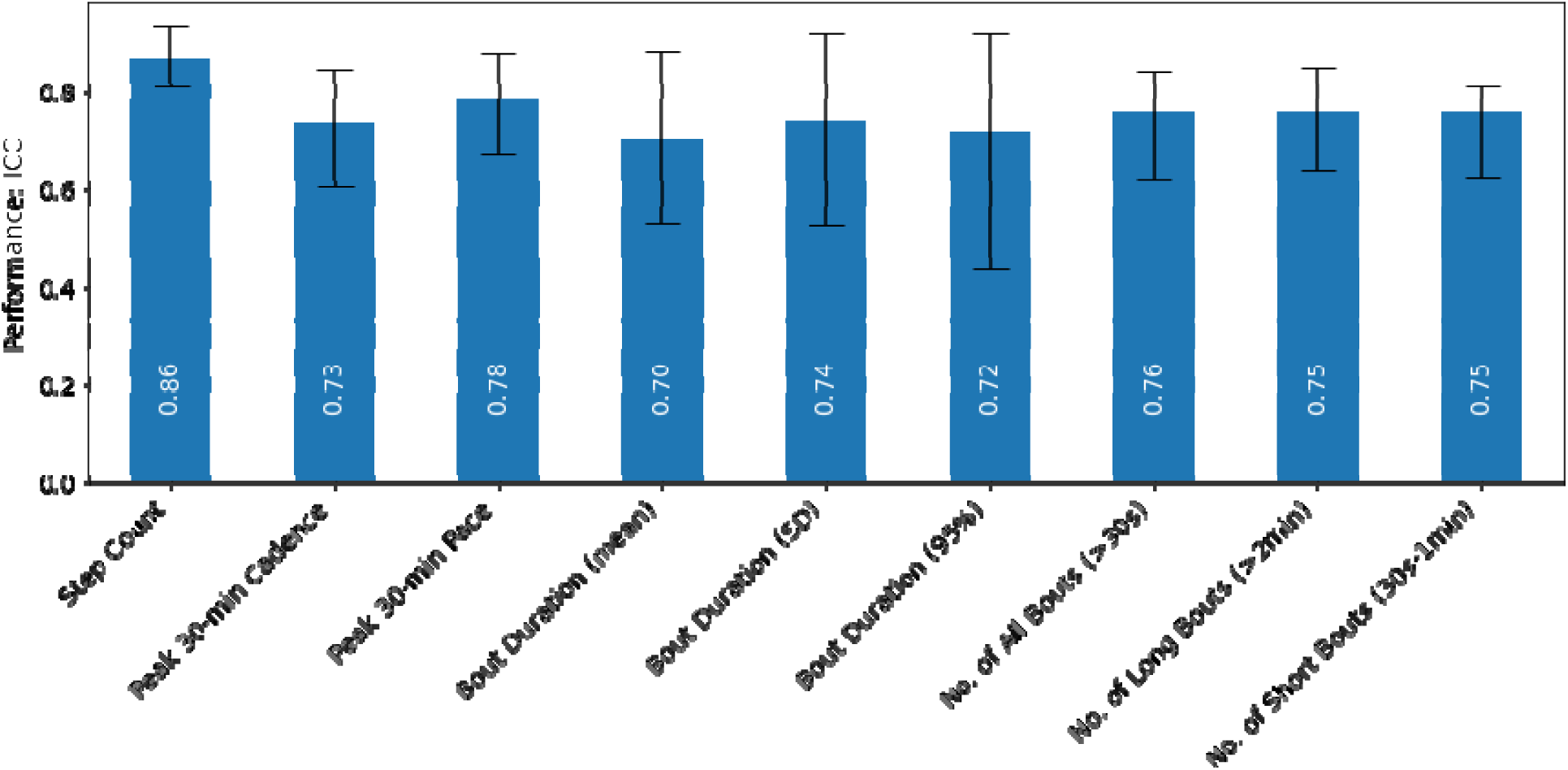
Accuracy Characterization: Intraclass Correlation Coefficient (ICC) results (and 95%CIs) obtained from the comparison of the digital measurements generated from the study device against those from the reference device. (Abbreviations: s=second; SD=standard deviation)

### Reliability characterization

For time-to-reliability characterization, the observation period ran for 20 months (April 13, 2018 - December 31, 2019). This analysis was designed to determine the duration of time (from 1 - 30 days) over which each measure needs to be aggregated (the different lengths of time tested were termed ‘aggregation scopes’) to yield stable values, indicating that it reliably captures an individual’s underlying behavioral state. Data were considered analyzable for this objective when participants had worn the device for at least double the duration of a given aggregation scope (in order to have data for 2 non-overlapping time windows), starting from a minimum of 2 days (for the shortest aggregation scope of 1 day) to a minimum of 60 days (for the longest aggregation scope of 30 days); in addition, at least 50% of the days in each time window had to have ≥ 12 hours of daily wear. The number of participants meeting these criteria varied according to the span of the aggregation scopes (N=234 for the one-day aggregation scope [that is, the smallest aggregation scope had the largest cohort]; n=81 for the 30-day aggregation scope [smallest cohort for the largest aggregation scope]) (Appendix Fig. 1b).

In this analysis, we included the same set of measures as for the accuracy characterization, except the 30-minute peak walking pace, since the measure is derived directly from 30-minute peak walking cadence (Table 1), therefore the results of this analysis were expected to be identical between these two measures. We calculated Fisher’s ICC’s between adjacent, non-overlapping windows of time for each aggregation scope (1 - 30 days). We computed a rolling mean for each daily aggregated measure over the set number of days for each aggregation scope, and then computed the ICC between adjacent windows. We repeated this computation by shifting the start date of each window by 1 day, and repeated the computation, testing aggregation scopes between 1 and 30 days.

## Results

### Accuracy Characterization

A total of 162 participant-days worth of data were collected from the 35 participants in the test cohort, with each participant contributing 1-10 days (median, 5). The mean daily step count, daily ambulatory time, and wear time per participant-day were 10,075.88 steps (standard deviation [SD], 4321.07),1.86 hours (SD, 0.78), and 13.73 (SD, 3.00) hours respectively (Appendix Fig. 2).

**Figure 2.**
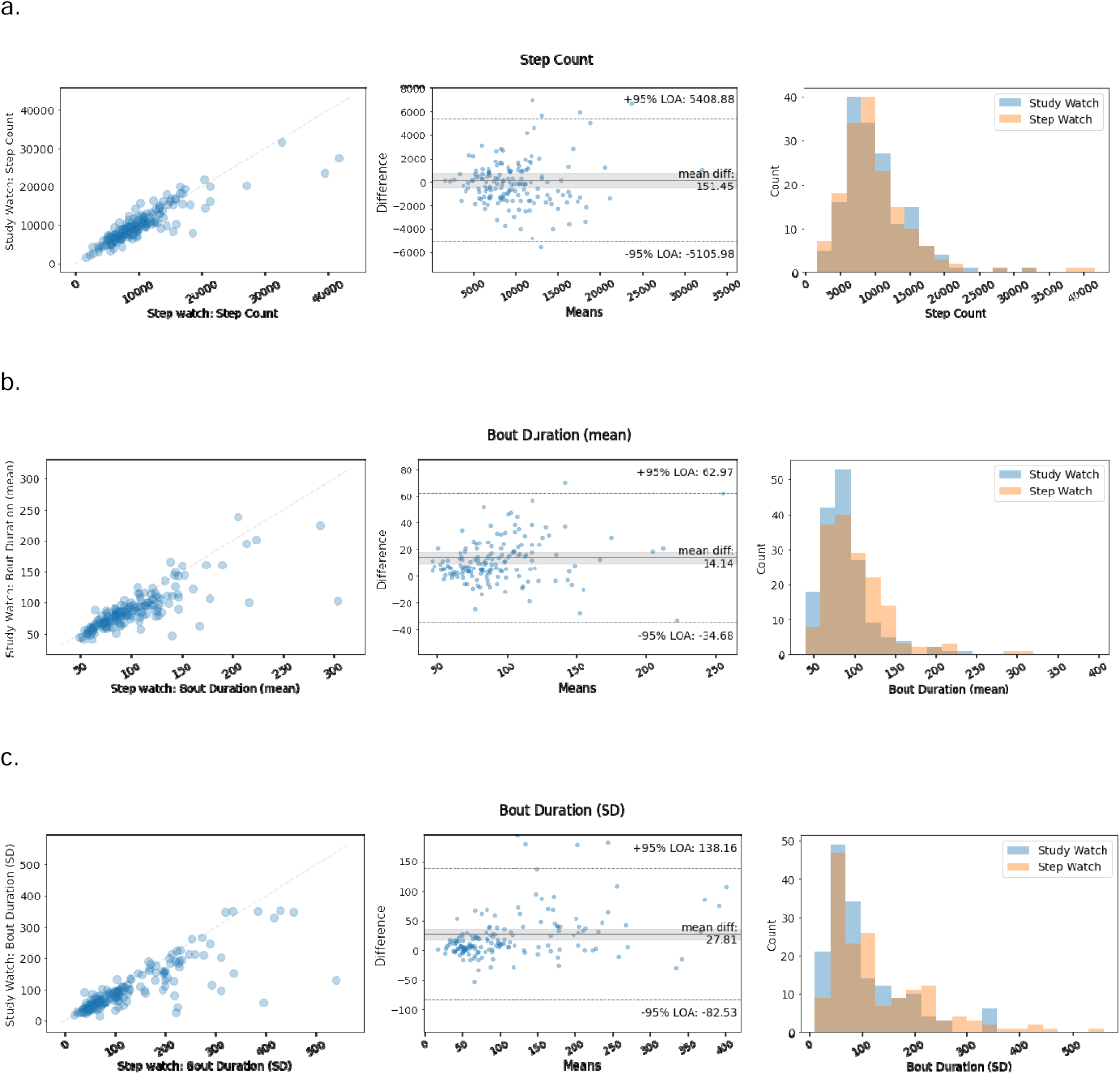
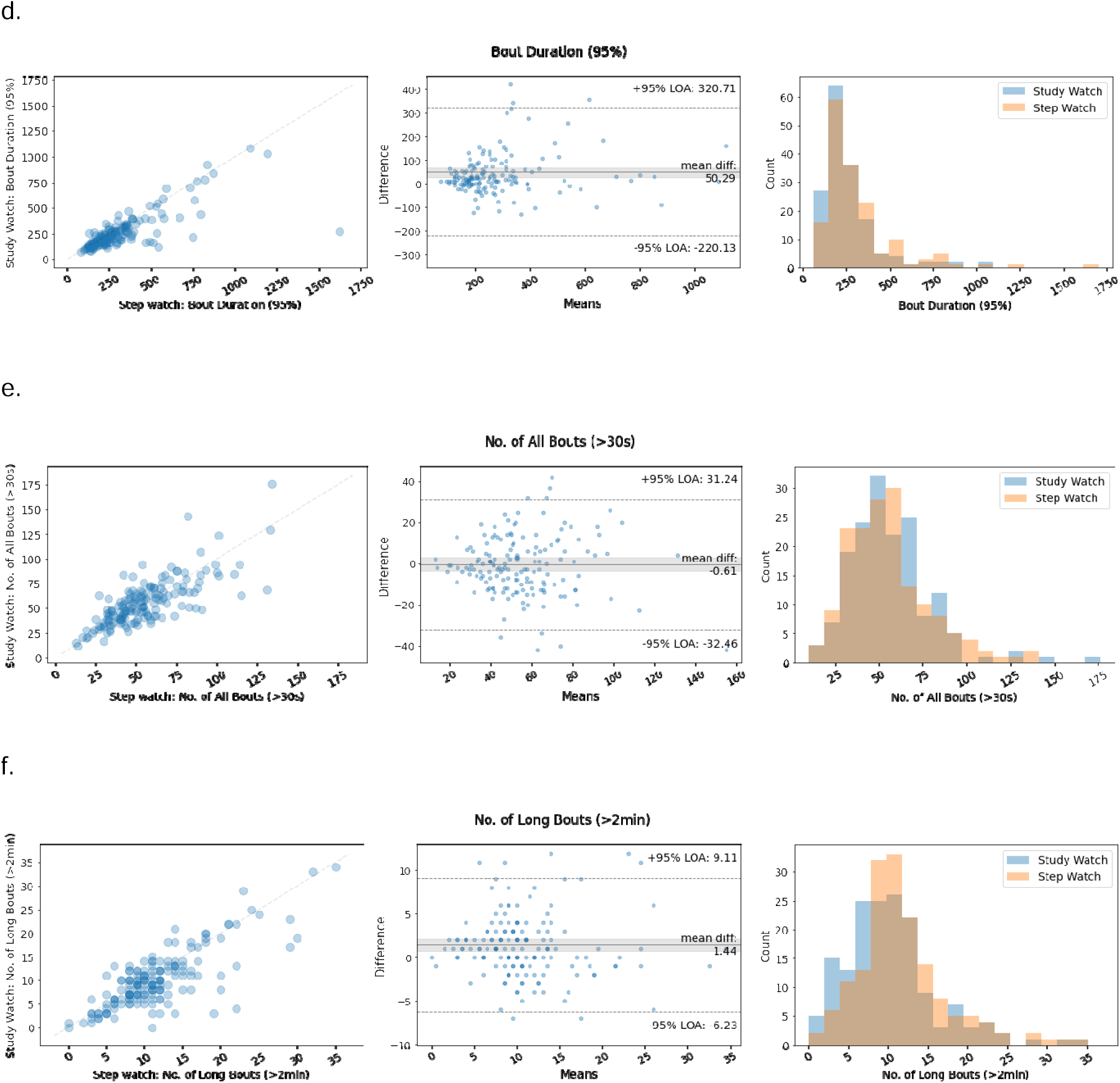
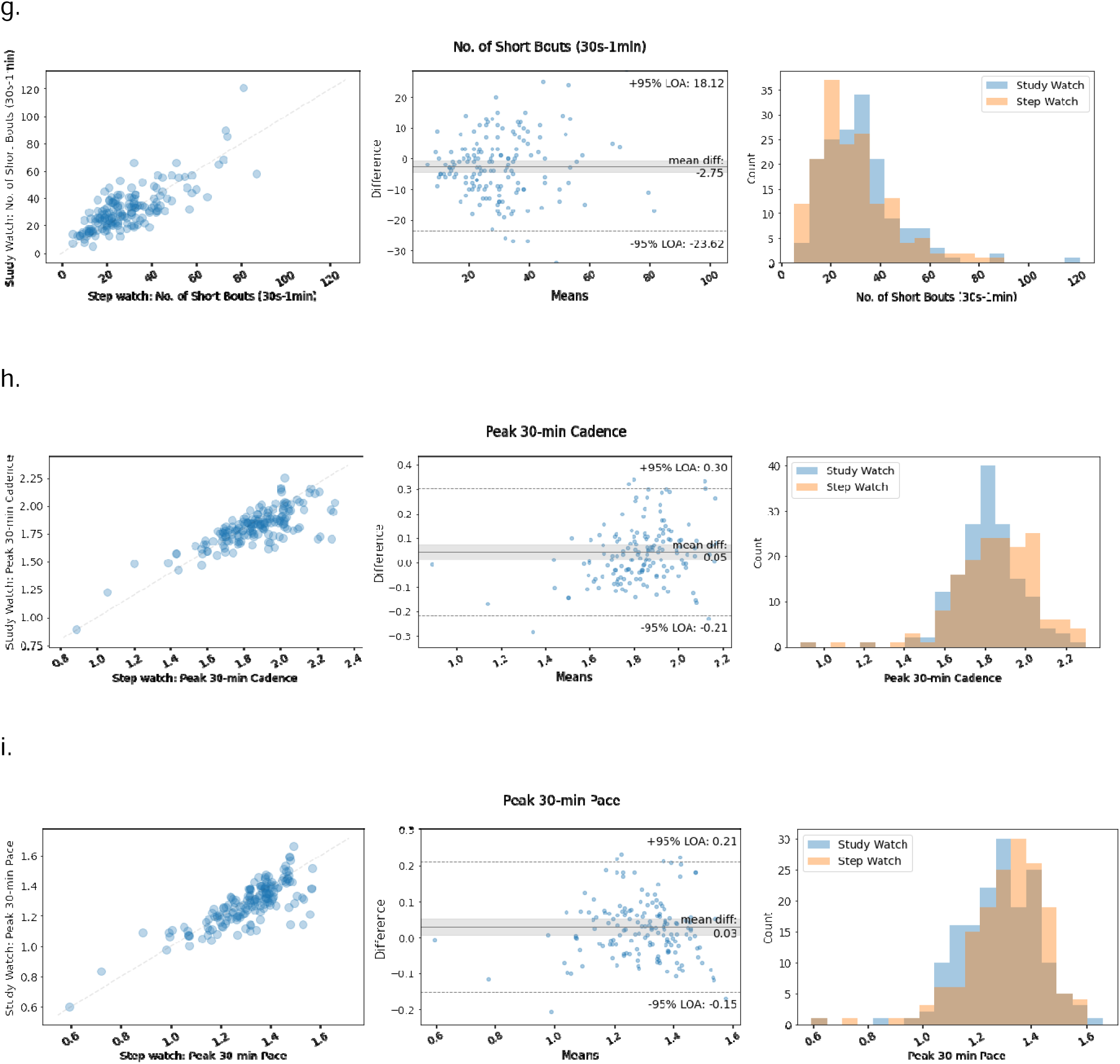
Accuracy Characterization: Detailed results of the comparisons of the digital measures generated from the study device against those from the reference device. Left column: plots of study device readouts (Y axis) vs reference device readouts (X axis). Middle column: Modified Bland-Altman plot showing the difference in mean values between devices (Y axis) vs mean values from the reference device (X axis). Right column: readout value distributions for both devices in the testing subcohort. (a) Daily step count. (b) Daily walking bout duration, mean. (c) Daily walking bout duration, standard deviation. (d) Daily walking bout duration, 95th percentile. (e) Number of daily walking bouts. (f) Number of daily long walking bouts. (g) Number of daily short walking bouts. (h) Daily peak 30-minute walking cadence. (i) Daily peak 30-minute walking pace. (Abbreviations: LoA=limits of agreement; s=second)

For each measure of interest (see Methods), the comparison of the values generated from the study device against the reference device showed ICC values ranging between 0.701-0.865 (Table 2, Figure 1); the measure ‘mean duration of daily bouts’ produced the lowest ICC value (0.701) and ‘daily step count’ had the highest (0.865). Pearson R values were all greater than 0.75 and all values were statistically significant (*P* <.001; Table 2). Bland-Altman analysis (Figure 2, middle) revealed that measure differences between the study and reference devices were not dependent on measure value without significant bias. Scatter plots (Figure 2, left) and distribution (Figure 2, right) of measures between study and reference devices showed overlap for all nine measures of walking.

**Table 2.**
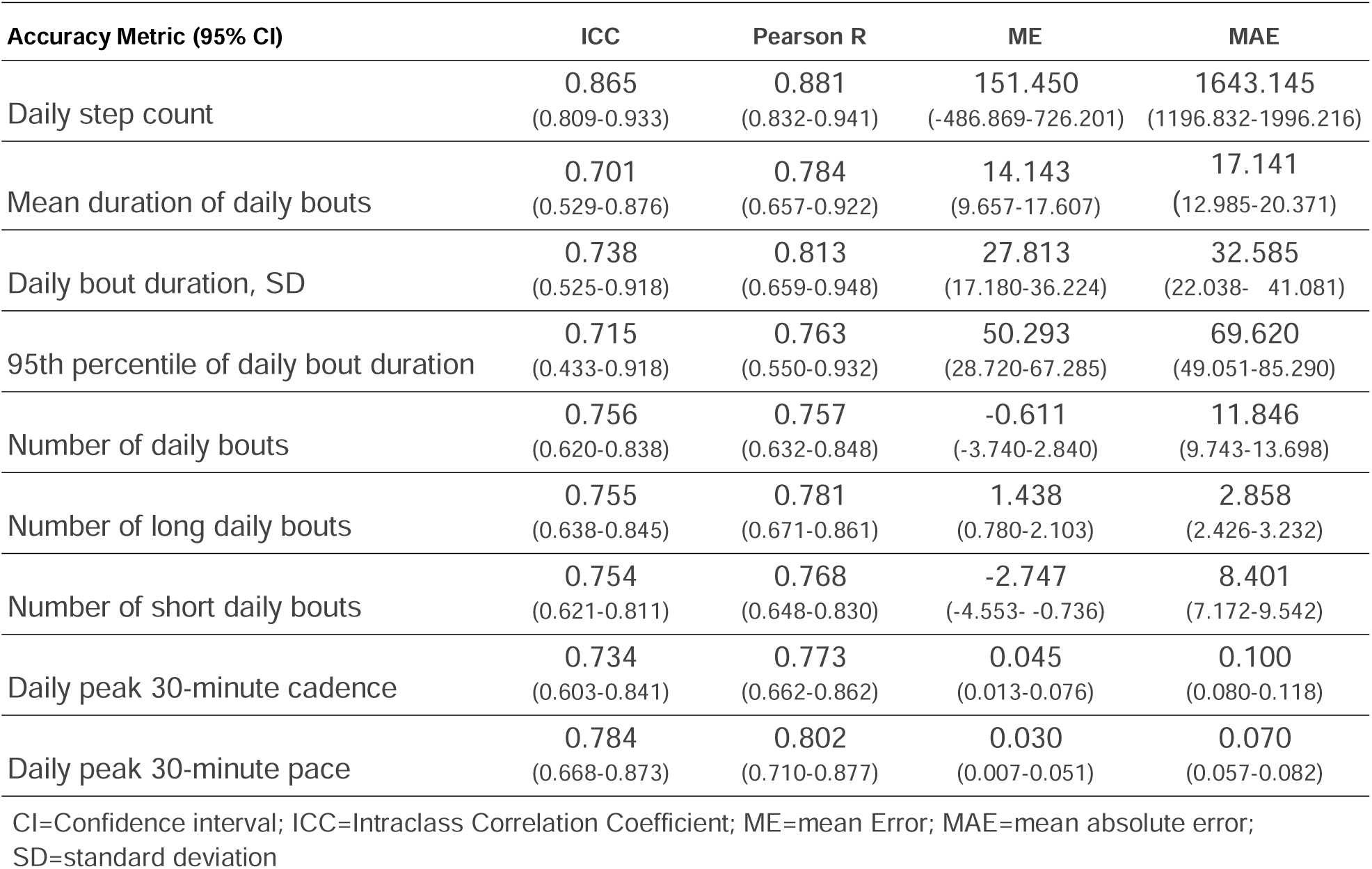
Summary of results from the characterization of accuracy of the measures generated from the study device, compared to those collected from the reference device (n=35).

### Reliability Characterization

In the cohort of eligible participants who yielded analyzable data (see Methods section, N=234), individual participant data were collected for up to 596 days (median, 119 days), for a total of 15,120 participant-days (see Appendix Fig. 1b). The mean daily step count, daily ambulatory time, and daily wear time per participant-day were 9701.06 (SD, 4321.88) steps, 77.42 (SD, 39.77) minutes, and 17.36 (SD, 4.04) hours, respectively.

We defined aggregation scopes of increasing duration from 1 day up to 30 days. For each of these scopes, the participant subcohorts that generated data deemed analyzable were of variable size (generally decreasing as the aggregation scope grew, Figure 3).

**Figure 3.**
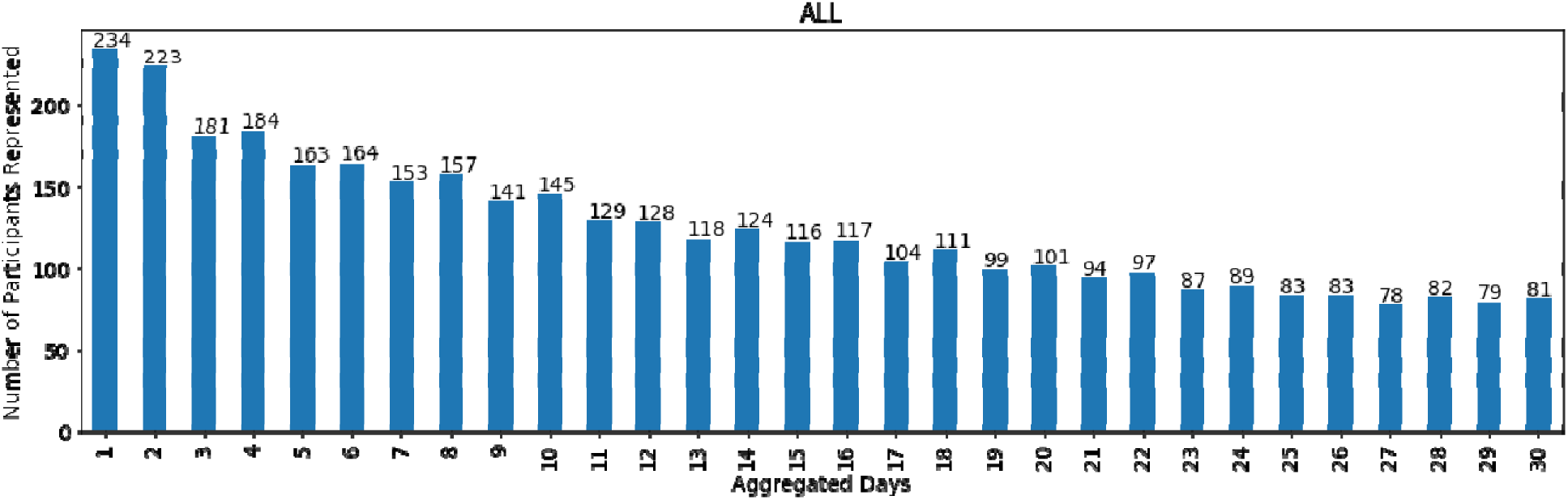
Time-to-reliability characterization: Size of the participant subcohorts with analyzable data across the aggregation scopes tested.

Across all the measures of interest in this analysis, the stability of the measure (estimated using ICC between adjacent time windows for readout) increased with longer aggregation scopes. The metrics ‘number of daily bouts,’ ‘bout duration, SD’ and ‘number of short bouts’ reached an ICC ≥ 0.75 at the earliest aggregation scope (12 days). Ultimately, all digital measures achieved an ICC ≥ 0.75 by 16 days, which we defined as the potential time-to-reliability benchmark in the context of this study (Figure 4). ICCs reached a plateau at values ranging between 0.78 and 0.84, depending on the measure.

**Figure 4.**
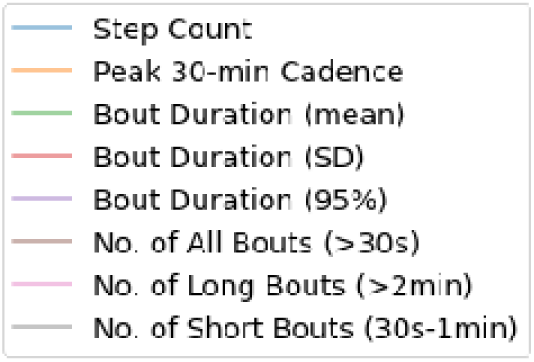
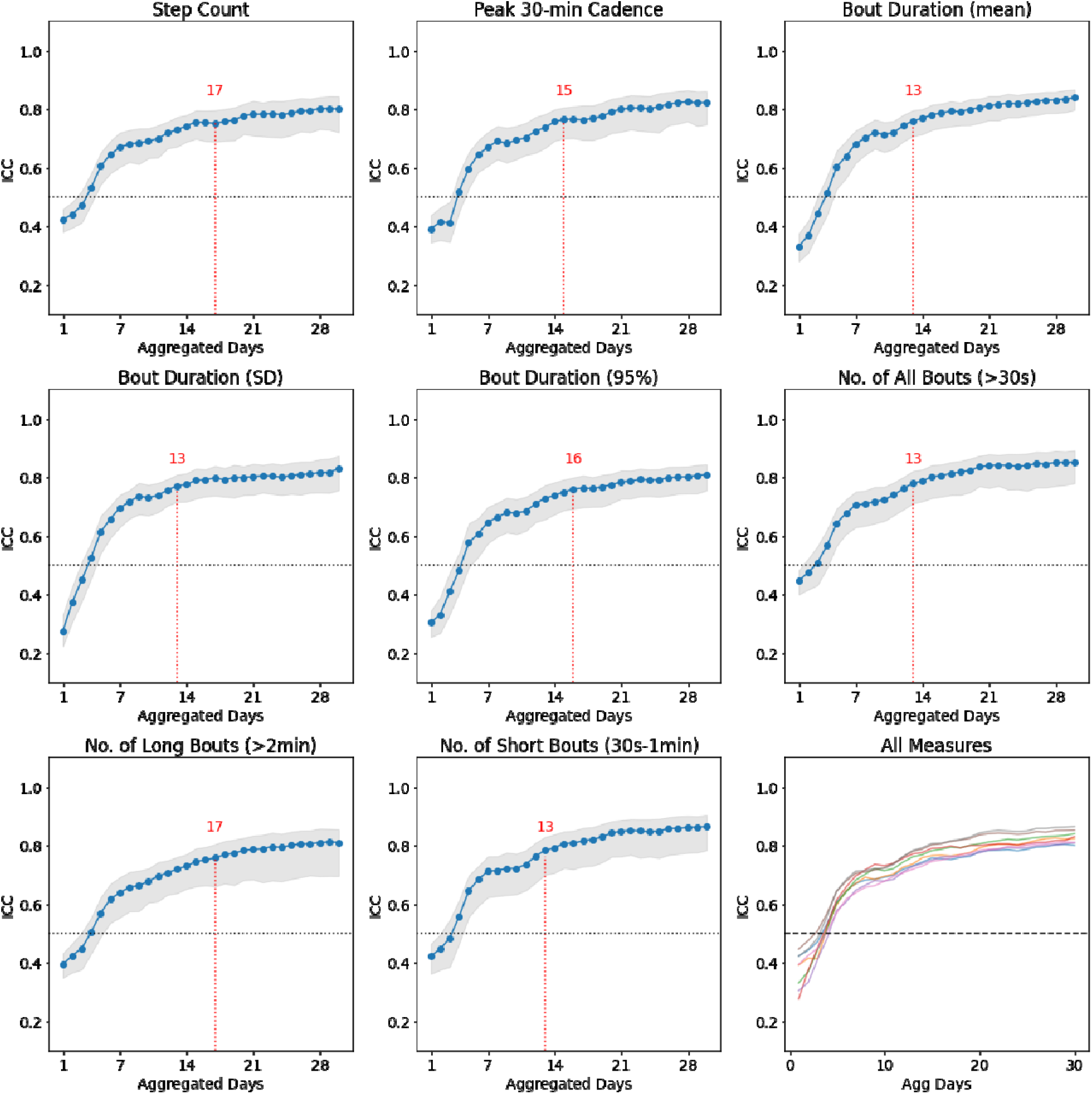
Time-to-reliability characterization: ICCs between adjacent readout windows according to aggregation scope duration, for the digital measures of interest. Line represents the ICC value plot, gray shading represents 95% CIs, and red annotations indicate the aggregation scope first exceeding an ICC value of 0.75. (Abbreviations: s=second; SD=standard deviation)

## Discussion

Mobility and walking behaviors represent meaningful aspects of health, known to be associated with quality of life in general and clinical prognosis in specific settings [33–35]. Therefore, improved methods to measure mobility and/or walking behaviors have the potential to improve clinical care and clinical trial efficiency. One of the goals of our research is to build accurate tools to objectively quantify aspects of walking behavior and extract clinically meaningful information in discrete populations of interest. In our prior work, we have developed algorithms whose outputs (measures for step counts and ambulatory time) demonstrated sensitivity to treatment effects in patients with Lewy Body Dementia [31]. We have also characterized the accuracy of an iteration of that algorithm in a cohort of diverse individuals in the real world [30]. This report expands upon that prior research, presenting a more comprehensive application of the algorithm that captures step count and other aspects of mobility, such as walking cadence and bouts. The objective of this study was to characterize the accuracy and reliability of this comprehensive set of digital measures of walking from users wearing a wrist-worn device in real world environments.

To our knowledge, this is the first report to characterize the amount of data required to ensure that a digital measure is reliable in a real-world setting. We developed a novel analytic approach to characterize time-to-reliability, that is, the time needed for a measure to reach a degree of stability. Time-to-reliability is an important consideration to inform the design of clinical studies tracking real world data, as it relates to specific metrics of interest. In this study, the time-to-reliability overall for all measures was less than or equal to 16 days (ICC ≥ 0.75 between non-adjacent readouts, for all measures at day 16, Fig. 4). This study included healthy individuals; in a clinical context, we anticipate that the stability of any given measure over time will be dependent on the type and severity of the disease of interest. It is reasonable to speculate that a mostly healthy cohort may demonstrate more variability and a larger distribution of walking behaviors than a cohort with disease burden, and this necessitates further research.

Most importantly, the algorithms developed to quantify daily step count and measures related to walking cadence and bouts were found to be accurate (agreement between the readouts from the study device and a highly-accurate reference device ranged between ICCs of 0.7-0.9 for all measures, Table 2). Our study approach captured measures of walking behavior in a real world setting, over multiple days, to closely resemble actual use cases. Based on these results, we believe that the performance demonstrated by these measures of walking supports their deployment in clinical trial settings with confidence.

Considering the exponential growth of research on wearable sensors and related devices in recent years, it is important to place the capabilities described in this report in that context. Our study approach included the use of a highly accurate but pragmatic, ankle-worn, ground truth label to assess multiple walking related measures in a real-world setting. Prior studies have used co-located investigational and reference sensors (e.g., two wrist-worn devices). Due to the known potential errors associated with body placement when capturing walking-related data [36–39], co-location could be vulnerable to bias towards overestimating performance, which our approach seeks to mitigate. Further, most studies have had a narrow focus on step counts [10–16], mostly in controlled laboratory environments and/or for limited time periods (e.g., single day in real-world setting). Thus, directly comparing study results side by side has to be done with caution, given the heterogeneity of the studies (comparisons of different devices, different ground truth sources, and with different analytic approaches), which highlights the need for standardization noted in professional statements in this field [6,9,40,41].

This study had limitations in regards to the participant population and the performance quality thresholds. First, our cohort was limited in size, and consisted of generally healthy participants. Future studies may be needed to characterize the generalizability of the performance of these measures in populations with particular kinetic hallmarks (e.g., neurological conditions, stroke, trauma) or with mobility capacity issues (e.g., cardiovascular or respiratory conditions). Our approach to determine time-to-reliability can be applied across studies in any therapeutic area and can guide study design requirements for wear-time compliance. One aspect that will require attention is the optimization of actual compliance with hypothetical protocol specifications about wear time, since this is a device intended for daily life use (for instance, our reliability analysis filtered participant-day data based on a threshold of 12 hours of daily wear time for 50% of the days over an evaluation period, but we did so retrospectively). Second, while we report on performance parameters, the definition of an acceptable performance (accuracy and reliability) quality threshold remains undefined in the field. We did not pre-specify performance categories in this study, but, for instance, prior accuracy studies have categorized agreement ICC values 0.7-0.9 as moderate-to-good [42,43], and reliability studies for patient-reported outcomes have considered test-retest ICC >0.5 as acceptable [44]. Importantly, what constitutes a clinically meaningful change for each of the measures of walking will likely depend on the therapeutic area under consideration. Further research is needed to address which of these measures can provide clinically-relevant insights in a given population. We believe that the accuracy and reliability results detailed here support the use of digital measures of walking as feasible and reliable endpoints in clinical studies.

In conclusion, we have developed algorithms that accurately quantify daily step count and measures of walking cadence and bouts from users wearing a wrist-worn device in a real-world setting. Further, we have also developed a novel method for characterizing the time required for a digital measure to stabilize (time-to-reliability). Given the growing use of wearable sensors to measure aspects of health, these findings may guide practical implementation of these digital measures of walking behavior into clinical studies.

## Disclosures/Acknowledgements page

### Data [/code/materials] sharing statement

Data from this study are not available due to the nature of this program. Participants did not consent for their data to be shared publicly.

### Authors’ contributions

Study concept and design: NK, ER, RK

Data collection: Verily Life Sciences LLC

Data analysis and interpretation: NK, SS, PB, SP

Draft writing and review: All

Draft approval for submission: All

## Acknowledgements

Authors wish to acknowledge writing and editing support from Julia Saiz from Verily Life Sciences.

## Abbreviations

ICC: Intraclass Correlation Coefficient (ICC)
ME: Mean Error
MAE: Mean Absolute Error
LoA: Limits of Agreement
SD: standard deviation (there are two contexts for this abbreviation in the report: as it refers to the variance in specific results of this analysis, or as it refers to one of the actual measures under study, i.e. standard deviation in the number of daily bouts)

## APPENDIX

### Step Count Algorithm for the Verily Study Watch

The underlying algorithm processes device user data in 10-second epochs. We trained a neural network on StepWatch generated ground-truth labels (n=35, same dataset as Pilot Program in Appendix Fig. 1). The following 14 features were extracted from the study device acceleration data, in 10-second epochs: 3 features related to deviations of the signal, 5 features derived from the power spectral density (PSD) energy in frequency bands typically associated with user’s ambulation (e.g., walking or running), 2 features that are signal percentiles (e.g. 95th percentile), and 4 features that are differences between signal percentiles (e.g. interquartile range). Detailed validation results of the ambulatory status classification algorithm were published previously [1].

**Appendix Figure 1.**
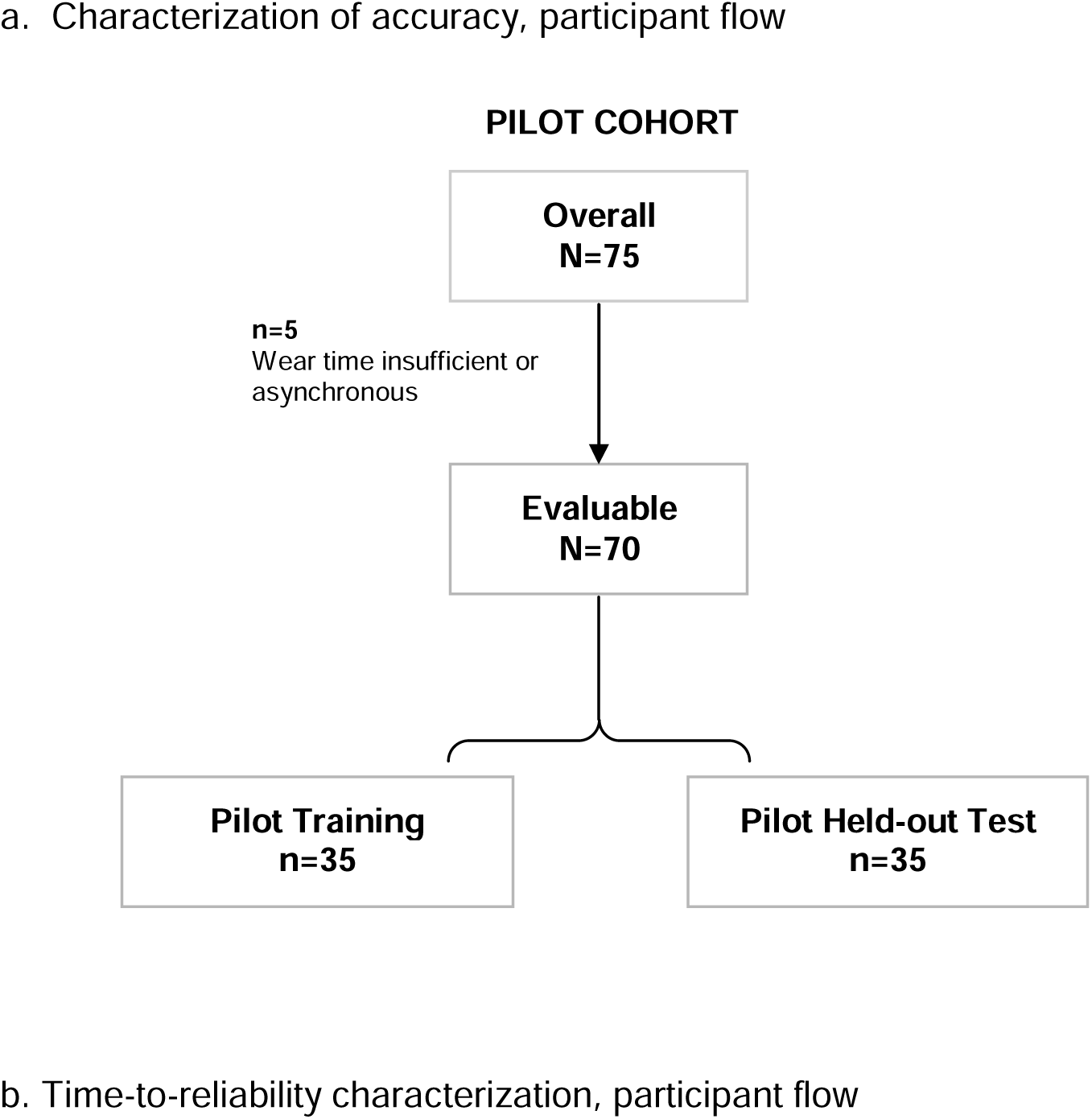
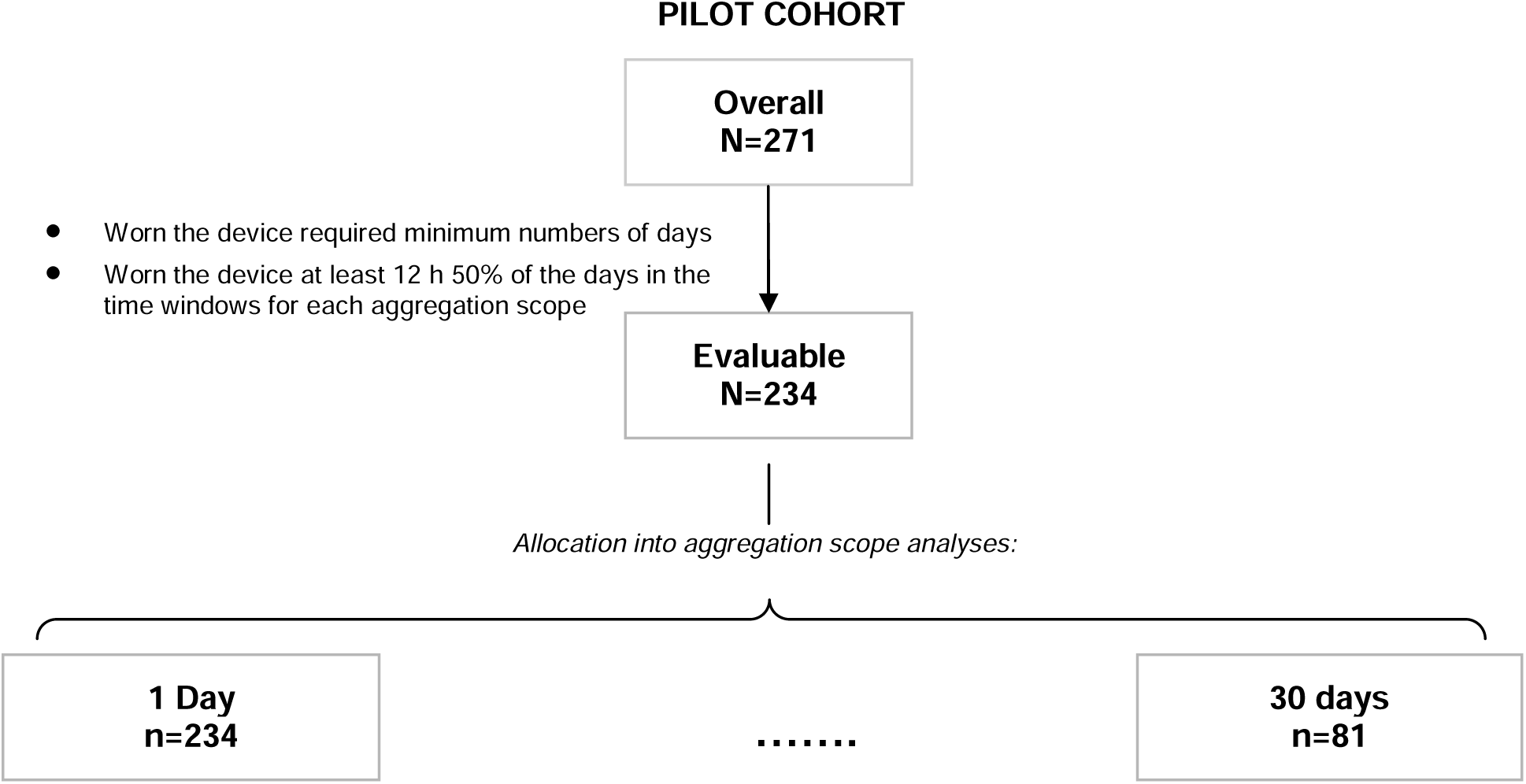
Participant flow into study cohorts/subcohorts

**Appendix Table 1.** Characteristics of the participants in the cohort for the accuracy characterization

|  |  | Accuracy Characterization Cohort<br>(N=75) |
| --- | --- | --- |
| Sex, n (%) | Female | 25 (35.7) |
|  | Male | 45 (64.3) |
| Age, yrs | Median | 30 |
|  | Mean (SD) | 33.2 (8.5) |
|  | Range | 23 - 63 |

**Appendix Figure 2.**
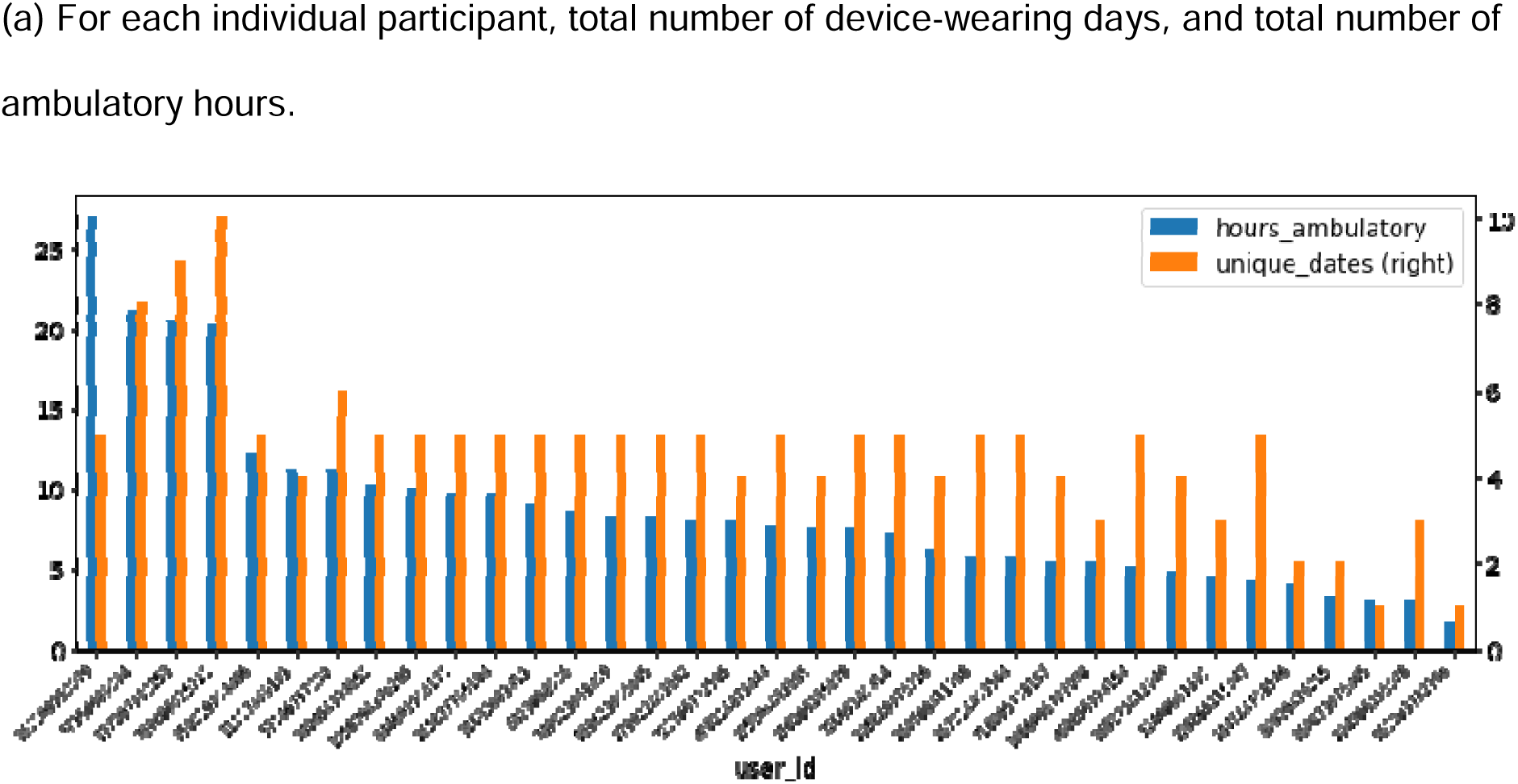
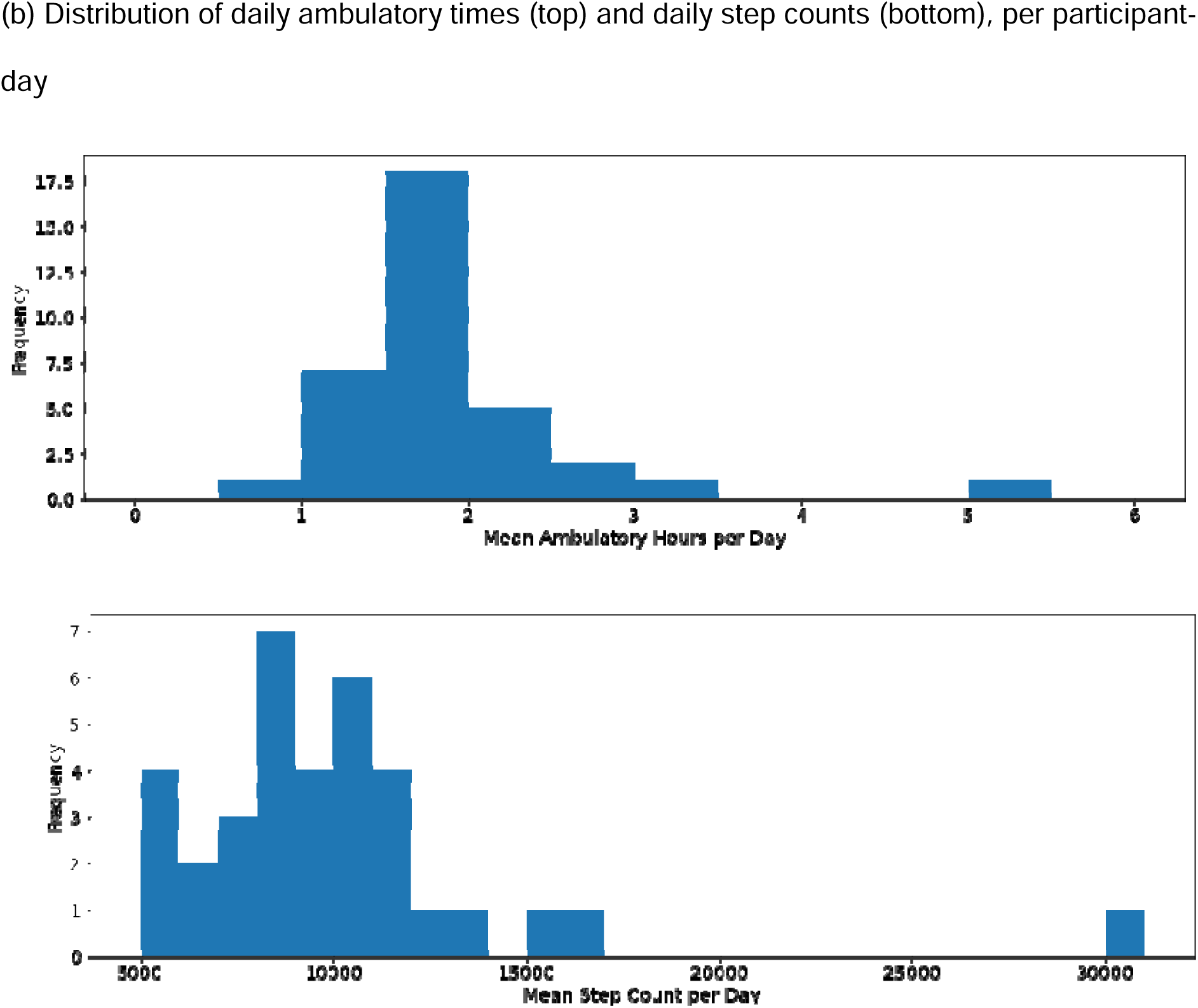
Characterization of accuracy: Overall features (in terms of distribution of walking activities) of the data collected from the testing subcohort.

